# 1,3- and 1,4-linked Polysaccharides Uptake in Intestinal Cells Relies on Clathrin/Dynamin 1/Rab5-dependent Endocytosis

**DOI:** 10.1101/2025.01.02.631151

**Authors:** Wenfeng Liao, Dianxiu Cao, Ying Wang, Zhenyun Du, Jian Yao, Pengfei Dou, Yuandong Zheng, Zhiming Wang, Xia Chen, Peipei Wang, Hao Chen, Xingxing Diao, Kaiping Wang, Kan Ding

## Abstract

Natural polysaccharides are considered to be a promising feedstock for manufacturing sustainable materials. However, polysaccharides are believed by most biologists that they could not be absorbed by intestinal mucosa to enter circulation system owing to their high hydrophilicity and large molecule size. Herein, β-1,3-linked glucan (GFPBW1) and α-1,4-linked glucan (WGE) is used as the model polysaccharide and the transport experiments show that different linked and charged polysaccharides can permeate Caco-2 cell monolayers. The *in vivo* experiments demonstrate that GFPBW1 and WGE are detectable in rodent plasma and liver after oral administration the isotope (^99m^Tc, ^3^H) and fluorescein labeled polysaccharides. Further study show that polysaccharide uptake requires clathrin heavy chain (CLTC) and its associated factors Rab5 and dynamin1 in intestinal cells using gene-knockdown strategies and inhibitors. Strikingly, polysaccharide absorption is attenuated in both *CLTC* intestine deficient mice and Rab5, dynamin1 conventional-knockout mice. Importantly, membrane receptors bone morphogenetic protein receptor type IA (BMPRIA), Dectin-1 and epidermal growth factor receptor (EGFR) are also critical for specific structural polysaccharide internalization. These findings provide novel insight to understand polysaccharide absorption mechanism and lay foundation for oral polysaccharide-based new drugs development.

## 1. Introduction

As one of the four fundamental biomolecules in nature, carbohydrates mediate various biological processes such as signal recognition, pathogenesis prevention, immune modulation and development.[1] Carbohydrate can be found as simple monosaccharide, oligosaccharide and polysaccharide or forming more complex glycoconjugates, such structural variability makes carbohydrate hot spots for biomedical intervention.[2,3] Early in 1940, carbohydrate-based antibiotics including streptomycin and gentamicin (both are aminoglycosides) were discovered as anti-infection drugs. Then, the ganglioside GM1 was developed for acute stroke treatment, hyaluronic acid and chondroitin sulfate (both are polysaccharide) were investigated as anti-arthritis drug.[4] Since 2000, tremendous progresses in glycoscience fields bring vast opportunities for novel carbohydrate-based drug discovery.[5] In 2019, the conditional launch of sodium oligomannate (GV-971) in China further dramatically boosts the innovative carbohydrate drug development to a new climax.[6] Surprisingly, polysaccharides represent the largest percentage of the recorded carbohydrate drugs.[7] To date, more than 40 commercial polysaccharides-based drugs have been launched including heparin, which is a high sulfated heterogenous glycosaminoglycan, was approved as an anticoagulant drug for more than 80 years.[8] Nevertheless, the biological roles of exogenous polysaccharides go further beyond due to their extensive therapeutic properties in antibiotic, antioxidant, anti-tumor, anticoagulant, antiaging and immuno- stimulation fields, etc.[9] Notably, about 17 approved or clinically developed polysaccharides-based drugs are administrated orally according to the records in Cortellis^TM^ (Drug Discovery Intelligence) database.[8] However, it’s largely unknown whether these bioactive polysaccharides may arrive at lesion site after oral administration. Thus, firstly, we are not sure whether the polysaccharides degradation occur or not after the polysaccharide uptake by intestine; Secondly, may those polysaccharides enter blood circulation system? More critically, there were few studies on polysaccharides absorption mechanism to date. The main reason is that most researchers believe polysaccharides cannot be absorbed by the gastrointestinal tract due to their large molecular size and high hydrophilicity. However, clinical studies demonstrated the absorption of exogenous high molecular mass chondroitin sulfate in the intestinal lining.[10] Nevertheless, it is challenge for clinically used heparin to pass through the gastrointestinal epithelium.[11] Hence, polysaccharides absorption by the intestinal lining is still a controversial topic.

Clearly, there is a critical need to understand polysaccharide pharmacokinetics to further fully utilize bioactive polysaccharides and develop novel polysaccharide-based drugs. However, owing to their large molecular weight and deficiency of UV-absorption groups, it is challenging to detect plasma polysaccharides which may be not distinguishable when the exogenous and endogenous polysaccharide consisting of same or similar monosaccharides. During the few last decades, many attempts have been made to address such challenge. The prior method is radioactive isotope labeling (such as ^14^C, ^125^I, ^35^S, ^3^H and ^99m^Tc), which were widely used in the pharmacokinetics investigations of polysaccharide.[12–17]Another common method used in pharmacokinetic studies is liquid chromatography coupled with tandem mass spectrometry (LC-MS/MS). However, the current chromatographic and mass spectrometric techniques have limited application due to large molecular size and lacking of chromophoric groups of polysaccharides. Luckily, there has been some progress with the development of spectrometric techniques, such as high-performance liquid chromatography (HPLC) followed by post-column fluorescence derivatization or pre-column fluorescein labeled.[18–19]

Except studying the ability of polysaccharides to permeate the intestinal mucosa, it is critical to study the absorption mechanism. Several pathways were reported involving in extracellular macromolecules uptake, such as micropinocytosis,[20,21]phagocytosis[22] and endocytosis.[23] Endocytosis is defined as the production of internal bodies from the plasma membrane and critical in antigen presentation, cell migration, and regulates many intracellular signaling cascades.[24] Eukaryotic cells exhibit at least two endocytic pathways: clathrin-dependent and clathrin-independent pathways. The mechanism of clathrin-mediated endocytosis (CME) is well studied.[25,26] The clathrin-independent pathways include caveolin-dependent and flotillin-dependent endocytosis, and phagocytosis.[27–29] Additionally, CME is required for the uptake of large molecules such as proteins, pathogenic bacteria and viruses in mammalian cells.[30,31] These knowledges inspire us to ask whether uptake of polysaccharide by intestine is clathrin dependent or clathrin-independent or something else.

In this study, we studied the ability of various polysaccharides to permeate intestinal cell layers *in vitro*. Moreover, GFPBW1 (β-1,3-linked glucan from *Grifola frodosa*[32]) and WGE (α-1,4-linked glucan from *Gastrodia elata* Bl[33]) as two model polysaccharides (two main types of polysaccharides in plant, fungus, and herbal medicine) were employed for the *in vivo* and mechanism study.

## 2. Results and Discussion

### 2.1 Different linked type and charged polysaccharides could be absorbed and internalized into intestinal cells *in vitro*

Caco-2 cell monolayer is widely used as an *in vitro* model mimic human small intestinal mucosa to predict orally administered drugs absorption.[34] To test the potential whether polysaccharide can be absorbed by intestine epithelial cells, transport studies for GFPBW1 and WGE were firstly performed. The results showed that the retention times of the transported GFPBW1 and WGE were not changed significantly compared with the prototypical polysaccharides in the HPLC chromatograms (Figure 1A), indicating the molecular sizes of the penetrated polysaccharides did not change much, further suggesting that polysaccharides might be transported by Caco-2 cell monolayer without significant degradation. This result surprised and encouraged us to ask whether this phenomenon is widely happened and further test the absorption situation of other linkage types and different charged polysaccharides. Hence, the transport experiments of lentinan (clinically used branched 1,3-linked glucan), MDG-1 (1,2-linked fructan), dextran (commercially available 1,6-linked glucan), GFPBW1-NH_3_ (amino-branched 1,3-linked glucan), WGE-NH_3_ (amino-branched 1,4-linked glucan), GFPBW1-SO_4_ (sulfated-branched 1,3-linked glucan) and WGE-SO_4_ (sulfated-branched 1,4-linked glucan) were then conducted. Surprisingly the results showed that diversely linked and differently charged polysaccharides could penetrate Caco-2 cell monolayers. In terms of the penetrated amounts, the value for GFPBW1 was close to that for dextran but significantly greater than that for WGE. Moreover, the penetrated amount of MDG-1 (with the smallest molecule size) was the largest, and was similar to that for lentinan (crude polysaccharide). Furthermore, the charged polysaccharides also displayed variation in the penetration values. The amount of GFPBW1-NH_3_ was dramatically more than that of GFPBW1 and GFPBW1-SO_4_. Oppositely, the amount of both WGE- NH_3_ and WGE-SO_4_ was higher than that of WGE (Figure 1B). Surprisingly, the apparent permeability coefficient (*P*_app_) values of the tested polysaccharides were more than 1 × 10^-6^ cm/s, indicating that they might be absorbed by the cells (Table S1, Supporting Information).

**Figure 1.**
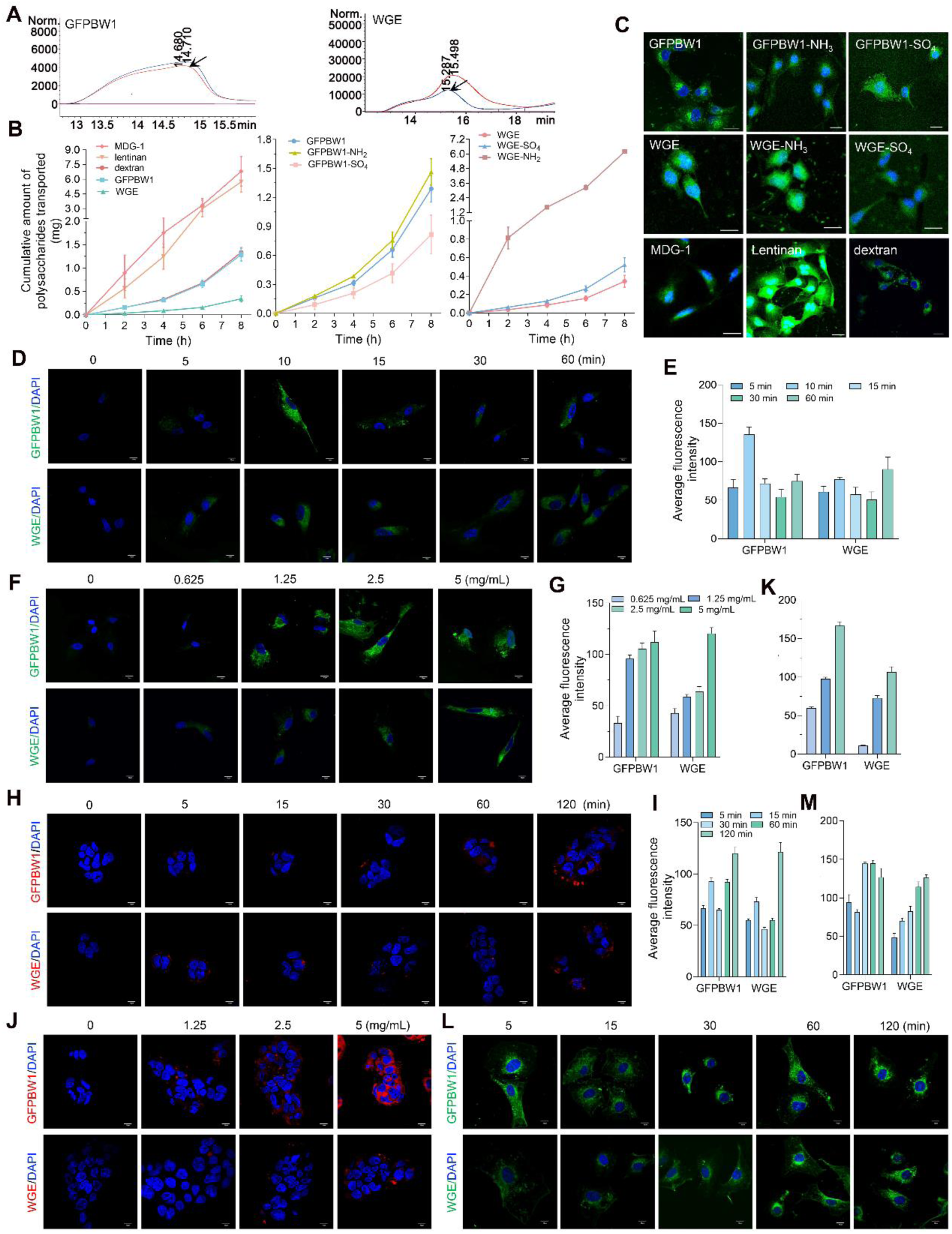
Different linked and charged polysaccharides could permeate Caco-2 cell monolayer and uptake by intestine cells. A) The transport experiments of GFPBW1 and WGE were performed as described in experimental section (Supporting Information). The chromatograms of the prototypical polysaccharides (arrowheads indicated) and samples (collected at 8 h) are shown. B) The Q-T curves of the polysaccharides with different linkages such as GFPBW1, WGE, dextran, MDG-1, lentinan; GFPBW1 and its derivatives (GFPBW1-NH_3_, GFPBW1-SO_4_); WGE and its derivatives (WGE-NH_3_ and WGE-SO_4_) are exhibited in the left, middle and right panel, respectively. C) Representative images of internalized polysaccharides in T24 cells. Scale bars, 20 μm. D-G) HIEC-6 cells were treated with fluorescein labeled GFPBW1 and WGE for the indicated time intervals (D) or at different concentrations (F), then cells were fixed and photographed by microscopy, mean fluorescence intensity was measured for all the fields of view for each group at three independent experiment and showed in (E) and (G), respectively. H-K) Similarly, images of the uptake experiments for GFPBW1 and WGE in Caco-2 cells in various time (H) or at different concentrations (J) are displayed, while (I) and (K) represent the quantified fluorescence intensity of (H) and (J). L-M) IEC-6 cells were treated with GFPBW1 and WGE at various time points and examined by confocal microscopy, measured fluorescence intensity is shown in (M), scare bars, 10 μm.

In order to trace polysaccharide transport, we fluorescein-labeled the polysaccharides,[35] the labeling positions was verified using raffinose, mannose via NMR and confirmed that the labeling procedure did not affect the structure and bioactivity of the saccharides too much (Figures S1-S4, Supporting Information). Next, the uptake experiment of different linkages and charged polysaccharides were performed. The results clearly showed that all the tested polysaccharides could be internalized into the cells (Figure 1C), while uptake of the fluorescein alone was not happened in the cells (Figures S5-S6, Supporting Information). Then, we asked whether time or/and concentrations will affect polysaccharides uptake. To better understand the details of polysaccharide penetration, GFPBW1 and WGE were employed as the model polysaccharides to do the following exploration. To address the question, human intestinal epithelial cell line (HIEC-6) was treated with GFPBW1 or WGE at different concentrations or for various time points. The results revealed that both GFPBW1 and WGE could be internalized into the cell as early as 5 min, both polysaccharides showed sharp increased internalization at 10 min, then the internalized polysaccharide decreased at 15 and 30 min, however the uptake of GFPBW1 was recovered in 1 h later (Figure 1D, E). Moreover, significant fluorescence could be observed at the concentration of 1.25 mg/mL of GFPBW1 and 2.5 mg/mL of WGE, respectively (Figure 1F). The uptake of both polysaccharides exhibited a concentration-dependent manner (Figure 1G). Subsequently, similar experiment was designed using Caco-2 cells. Similarly, GFPBW1 and WGE showed analogous uptake time dynamics in this cell line (Figure 1H, I), and both demonstrated a concentration dependent uptake manner (Figure 1J, K). Furthermore, polysaccharides uptake time dynamics was examined in IEC-6 (rat intestinal cell line), WGE showed a time-dependent pattern. Conversely, GFPBW1 showed a decrease in 15 min and recovered after 30 min (Figure 1L, M). Thus, the uptake dynamics of GPFBW1 and WGE in rat and human cell lines was different. The reason of such difference between the two polysaccharides might be partially due to their structures.

### 2.2 Polysaccharides were absorbed in rat and mice intestinal lining *in vivo* and could be detected in rodent blood and liver

Next, we sought to ascertain whether the two model polysaccharides could be absorbed *in vivo*. GFPBW1 and WGE were orally and intravenously administered to rats, while commercially available lentinan was also included as a control. Firstly, the spectrum of blank rat plasma mixed with GFPW (internal standard) or without followed extraction as described in supplemental methods were displayed (Figure S7, Supporting Information). The HPLC chromatograms showed the retention time and peak pattern of lentinan before and after the pretreatment procedure did not change obviously, indicating good efficiency of the extraction process (Figure S8, Supporting Information). Surprisingly, GFPBW1 was quantifiable in all plasma samples following oral and intravenous administration (Figure 2A, upper panel). Similar results were obtained for WGE and lentinan, and there were trace changes between the retention time and peak pattern of the plasma polysaccharides and their prototypical ones (Figure 2A; Figure S9, Supporting Information), suggesting that the tested polysaccharides were absorbed at least partially as macromolecular form with little degradation *in vivo*. The pharmacokinetic profiles and parameters of GFPBW1, WGE and lentinan by oral administration or intravenous injection are shown in Figures S10-S12 and Table S2, 3, respectively.

**Figure 2.**
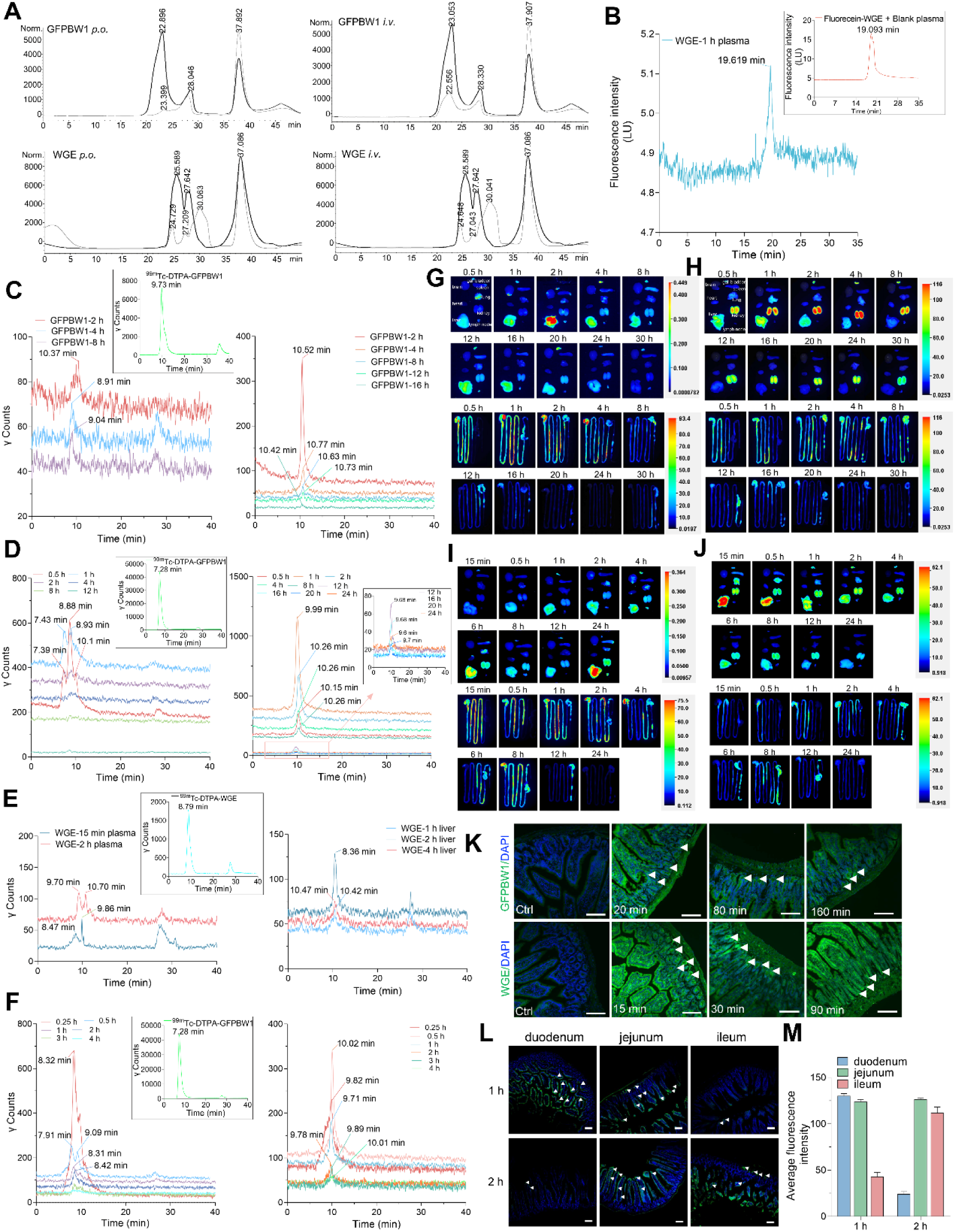
Polysaccharides could be absorbed by the rat and mice intestine *in vivo* and could be detected in rat and mice plasma and liver. A) The chromatograms of GFPBW1 and WGE before and after oral administration or intravenous injection in rat are shown. The solid and thin lines represent the prototypical or plasma polysaccharides, respectively. B) The HPLC chromatograms of fluorescein labeled WGE mixed with blank C57BL/6J mice plasma (embedded panel) and plasma after oral administration to mice are shown. (C-D) Moreover, the Radio HPGPC chromatograms of plasma (left panel) and liver lysate (right panel) after oral administration (C) or intravenous injection (D) of ^99m^Tc-GFPBW1 to C57BL/6J mice for the indicated time intervals were exhibited, the embedded photograph was the Radio HPGPC chromatograms of the original ^99m^Tc-GFPBW1. (E-F) Meanwhile, C57BL/6J mice were also oral administrated or intravenous injection with ^99m^Tc-WGE, and the Radio HPGPC chromatograms of plasma (left panel) and liver lysate (right panel) were shown in (E) and (F), the chromatogram of original radio labeled WGE was exhibited in the embedded picture. (G-J) The images of isolated organs (upper panels) and the whole intestine (lower panels) after the BALB/c mice were oral administrated (G for GFPBW1, I for WGE) or intravenous injection (H for GFPBW1, J for WGE) of Cy5.5 labeled GFPBW1 or WGE for the indicated time intervals. (K) Frozen section images of the intestinal tissues were shown after BALB/c mice were orally administered with fluorescein labeled GFPBW1 and WGE for the indicated time points. Scale bars, 100 μm. (L) Similarly, images of duodenum, jejunum and ileum of C57BL/6J mice after orally administrated with fluorescein labeled WGE for 1 h or 2 h were presented, white arrowheads indicate the internalized polysaccharides. The mean fluorescence intensity of each group was quantified in (M).

To further validate these findings, fluorescein-labeled GFPBW1 and WGE were labeled and detected on HPLC with a fluorescence detector. The results showed that both GFPBW1 and WGE demonstrating a typical peak pattern with measurable signals (Figure S13, Supporting Information). Next, C57BL/6J mice was orally administrated by fluorescein labeled WGE, followed by the polysaccharide extraction in mice blood and HPLC detection. The results showed that a typical peak of WGE from the plasma sample was presented on HPLC and the retention time was very closed to the original fluorescein labeled WGE after orally administrated for 1 h (Figure 2B), indicating that WGE could be absorbed by mice as macromolecule form.

To more precisely detect the polysaccharide after the oral administration in mice, we next sought to label the polysaccharides by radioactive ^3^H with the help of QZ Isotech (Shanghai, China) and examined by HPLC connected with beta ray detector. Indeed, chromatograms showed a typical peak with quantitative signals (Figure S14, Supporting Information). Subsequently, C57BL/6J mice were orally administrated by ^3^H-GFPBW1 and WGE, and the concentration-time curves of GFPBW1 and WGE in mice serum are displayed in Figure S15, further confirmed the results we obtained from rats and fluorescein labeled polysaccharides in mice. Further, we were still surprised to observe a peak with quantitative signals in mice plasma after oral administration of both ^99m^Tc labeled GFPBW1 and WGE (Figure 2C, E, left panels). More importantly, we even obtained canonical peak in mice liver lysate after oral administration of both GFPBW1 and WGE, and the retention time of liver lysate was closed to the original labeled ones, further implied that polysaccharides were absorbed partially as original macromolecular form without significantly degradation (similar molecular weights) within the test time and could accumulate in the liver (Figure 2C, E, right panels). Moreover, we obtained similar results from the plasma and liver lysate of mice after intravenously injected ^99m^Tc labeled GFPBW1 and WGE (Figure 2D, F). The pharmacokinetic parameters of ^99m^Tc- GFPBW1 and WGE in mice were summarized in Table S4, 5, respectively. Furthermore, fluorescence labeled GFPBW1(Figure 2G, H) and WGE (Figure 2I, J) with Cy5.5 were employed to trace the polysaccharides *in vivo* using living imaging after both orally administration (Figure 2G, I) and intravenous injection (Figure 2H, J). Obvious fluorescence was observed in both mice liver and intestine, further indicating the absorption of both polysaccharides.

Next, to test when the polysaccharides start to be absorbed by mice intestine, BALB/c mice were orally administered with fluorescein-labeled GFPBW1 and WGE, and we indeed visualized the internalized GFPBW1 and WGE in mice intestine after the oral administration for 20 and 15 min, respectively. However, the maximum amounts of internalized GFPBW1 and WGE occurred after the oral administration for 80 and 30 min, respectively (Figure 2K). In order to rule out the possible interference caused by mouse strain, we performed similar experiment in C57BL/6J mice. Microscopic analysis showed that polysaccharides internalized mostly into the intestinal enterocyte of the duodenum segments in WGE-1 h group, while there was less and trace internalized WGE in the jejunum and ileum segments. Conversely, we observed different absorption pattern in WGE-2 h group. The internalized WGE was increased sequentially in duodenum, jejunum and ileum segments, and even could be observed in mice ileum crypt (Figure 2L, M). Collectively, we demonstrated that polysaccharides could be absorbed by intestinal cells *in vivo* and detectable in rodent blood and liver.

### 2.3 Clathrin mediated endocytosis is critical for polysaccharides uptake in cultured cells

Some macromolecules such as EGFR and asialoglycoproteins were reported to be transported into the cells via endocytosis.[36,37]Inspired by this, we speculated that polysaccharides, which are hydrophilic macromolecules, might be absorbed via vesicular endocytosis. Firstly, we confirmed the localization of internalized polysaccharides using DiI to label cell membrane, the internalized polysaccharides were observed mostly in cytoplasm and partially on cell membrane (Figure 3A).

**Figure 3.**
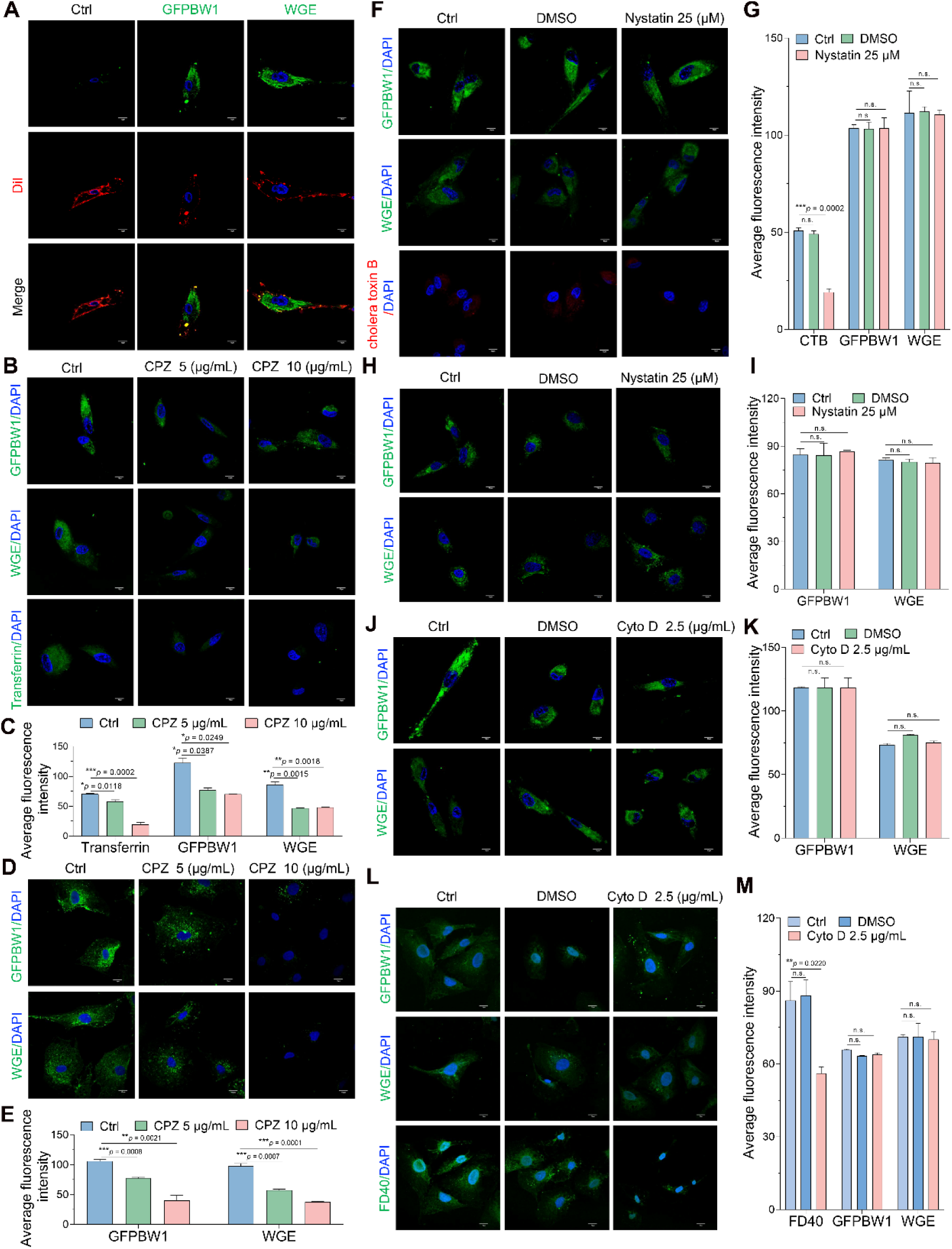
Disruption of Clathrin-mediated endocytosis attenuated the uptake of polysaccharides. A) HIEC-6 cells were incubated with GFPBW1 or WGE for the indicated time, then cells were fixed and stained with DiI (red). Subsequently, HIEC-6 cells were pretreated with different inhibitors and the uptake experiments of GFPBW1 and WGE were then performed, the images are displayed in (B), (F) and (J), separately. Scale bars, 10 μm. The mean fluorescence intensity was quantified in (C), (G) and (K), respectively. Similarly, the uptake experiments for GFPBW1 and WGE in IEC-6 cells were conducted after treating with CPZ (D), Nystatin (H) or Cyto D (L). Scale bars, 10 μm. The quantified results were exhibited in (E), (I) and (M), respectively. Data are represented as mean ± SEM.* *p* < 0.05, \*\**p* < 0.01, \*\*\**p* < 0.001 by unpaired Student’s *t* test (transferrin, cholera toxin B, and FD40 were marker of clathrin-mediated endocytosis, lipid raft mediated endocytosis, and micropinocytosis, fluid-phase endocytosis, respectively).

Next, we utilized inhibitors to block various endocytosis routes and then measured polysaccharides uptake in intestinal cells. HIEC-6 cells were pretreated with the chloropromazine (CPZ, inhibitor of CME), Nystatin (inhibitor of lipid raft mediated endocytosis) or cytochalasin D (Cyto D, inhibitor of micropinocytosis) for indicated time, then incubated with polysaccharides or endocytosis route markers. Interestingly, the uptake of both GFPBW1 and WGE was dramatically inhibited after CPZ disturbation, while transferrin was used as the marker of CME (Figure 3B, C), suggesting that polysaccharide internalization might be mediated by CME. Then, similar studies were also performed on IEC-6 cells. Consistently, uptake of GFPBW1 and WGE were reduced in CPZ treated group than that of control cells (Ctrl) in IEC-6 (Figure 3D, E). However, the uptake of both GFPBW1 and WGE in Nystatin treated HIEC-6 cells showed no obvious difference with that of Ctrl and DMSO group, while the uptake of cholera toxin B (marker of lipid raft mediated endocytosis) was nearly abolished (Figure 3F, G), which indicated the effective inhibition of lipid raft endocytosis. We observed similar phenomenon in IEC-6 cells (Figure 3H, I). Furthermore, the fluorescence images indicated that the internalization of GFPBW1 or WGE was not affected by Cyto D (Figure 3J, K), this observation was confirmed in IEC-6 cells under the same condition while the uptake of FD40 (marker of macropinocytosis or fluid-phase endocytosis) was severely impaired by Cyto D (Figure 3L, M). The above data suggested that internalization of GFPBW1 or WGE might be mediated by CME but not through macropinocytosis or lipid raft mediated endocytosis. It is reported that cell membrane-expressed proteoglycan is internalized via caveolin- containing endosomes.[38] Thus, we tried to determine whether caveolae-mediated endocytosis was involved in exogenous polysaccharides internalization. A caveolin- free Huh-7 cell[39] was therefore used to test the hypothesis. However, out of our expectation, the tested polysaccharides could still internalize into Huh-7 cells, further implying that polysaccharides uptake did not relay on caveolar lipid raft endocytosis (Figure S16, Supporting Information).

### 2.4 Polysaccharides uptake was impaired in *CLTC* knockdown intestine cells and *CLTC* deficient animals

To identify critical proteins during polysaccharide internalization process, HIEC-6 and IEC-6 cells were treated with fluorescein-labeled GFPBW1 and WGE, followed by immunostaining with anti-clathrin heavy chain (CLTC). Intriguingly, we found that the internalized polysaccharides could co-localize with CLTC protein in both HIEC-6 and IEC-6 (Figure 4A-D), indicating that CLTC might play a direct role in polysaccharides uptake. More importantly, we further observed the co-localization of the internalized polysaccharides and CLTC protein in Huh7 cells (Figure S16, Supporting Information). To further ascertain that CLTC is required for polysaccharide uptake, *CLTC* was knockdown in HIEC-6 cells in the presence of both GFPBW1 and WGE. The results showed that the uptake of both GFPBW1 and WGE were significantly decreased in the *CLTC* knockdown cells, while the efficiency of siRNAs was confirmed by immunoblotting (Figure 5A-C). This result encourages us to ask whether this situation also occur in animals. Then, we generated IEC-specific *CLTC*-knockout mice using Cre-loxP conditional gene targeting CLTC (*CLTC* CKO mice). The fluorescein labeled WGE was orally administrated to both the wild type and *CLTC* CKO mice. Surprisingly, we observed trace amount of internalized WGE in the ileum of the *CLTC* CKO mice, while obvious fluorescence was observed in the ileum of wild type mice, suggesting that silencing *CLTC* sharply blocked the uptake of polysaccharides *in vivo* (Figure 5J, K). Hence, our findings revealed a fundamental role of CLTC in polysaccharide uptake with these *in vitro* and *in vivo* validations.

**Figure 4.**
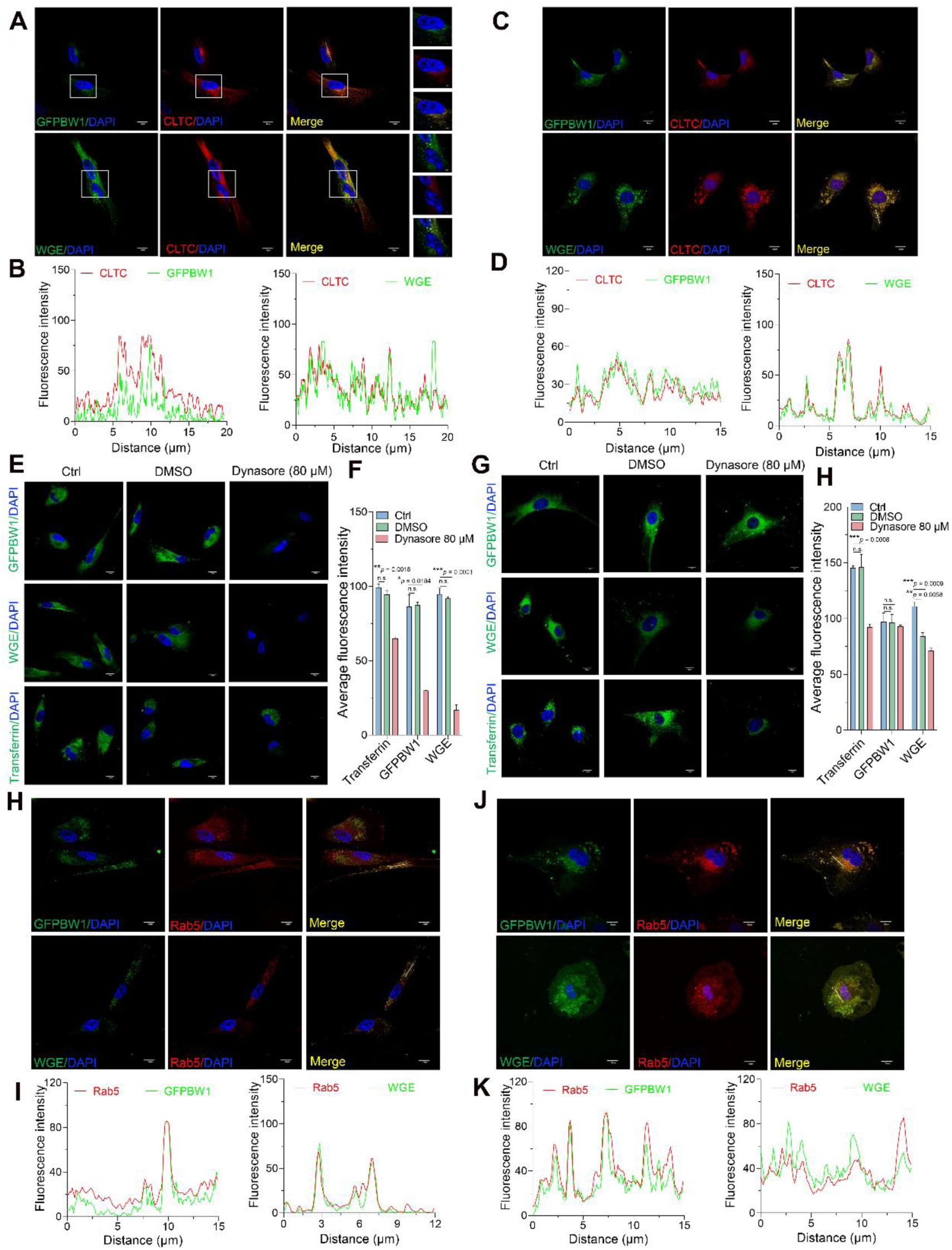
Clathrin heavy chain, dynamin 1 and Rab5 were required for the internalization of polysaccharides. HIEC-6 (A) or IEC-6 cells (C) were treated with fluorescein labeled polysaccharides and then subjected to immunostaining with clathrin heavy chain (CLTC). Scare bars, 10 μm, 2 μm for the enlarged images. The colocalization analysis results are showed in (B) and (D), respectively. Then, HIEC-6 (E) or IEC-6 cells (F) were treated with dynasore (dynamin inhibitor) and then performed the uptake experiment of GFPBW1 or WGE. Scare bars, 10 μm. The intensity was quantified in (F) and (H), respectively, \*\**p* < 0.01, \*\*\**p* < 0.001. Further, images of fluorescein labeled polysaccharides and immunostaining with anti-Rab5 in HIEC-6 (H) and IEC-6 cells (J) are displayed, colocalization was analyzed and showed in (I), (K).

**Figure 5.**
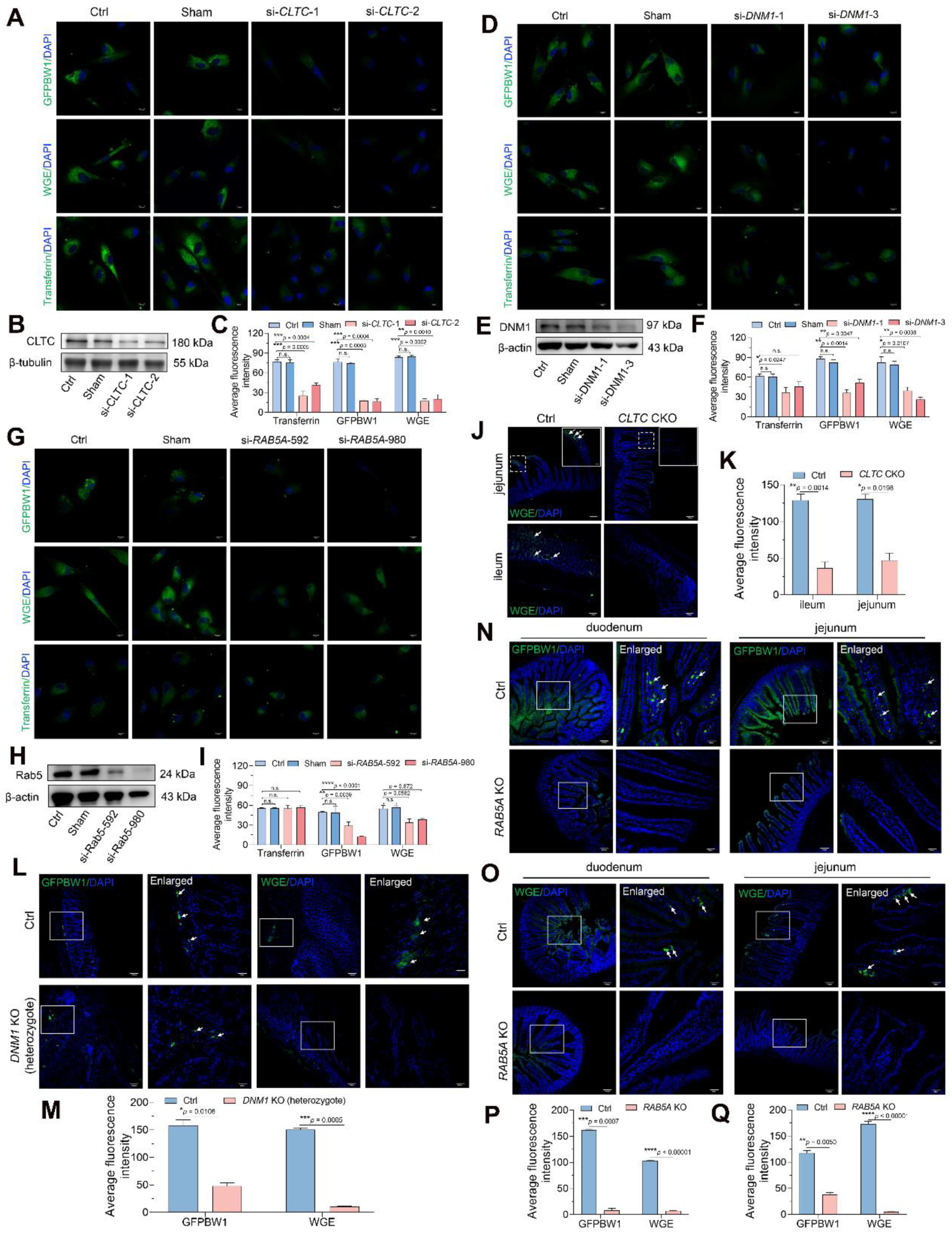
Internalization of polysaccharides was impaired by knockdown of clathrin heavy chain, dynamin 1 or Rab 5 *in vitro* and knockout of these proteins *in vivo*. HIEC-6 cells were transfected with si-*CLTC* or scrambled siRNA (Sham), the uptake experiments results are displayed in (A), siRNA efficiency was detected by Western blotting (B), the fluorescence intensity was measured in (C). Further, images of the internalized GFPBW1 and WGE in HIEC-6 cells after the disturbing of si-*DNM1* are presented in (D). Western blotting results showed the siRNA efficiency (E), the quantified results are shown in (F). Meanwhile, uptake experiments for GFPBW1 and WGE were conducted after the transfection with si-*RAB5A*, images are shown in (G), the immunoblotting and the fluorescence intensity results are presented in (H) and (I), scare bars, 10 μm. (J) Representative images of the jejunum and ileum from the *CLTC*^flox/flox^ (Ctrl) and *CLTC* CKO mice after oral administration of fluorescein labeled WGE, scare bars, 100 μm, the mean intensity was analyzed and showed in (K). (L) Images of the internalized polysaccharides in mice intestine from wild type mice (Ctrl, upper panel) and *DNM1 KO* mice (heterozygotes, bottom panel), the intensity was quantified in (M). Control and *RAB5A* KO mice were oral administrated with fluorescein labeled GFPBW1 (N) and WGE (O), images of the duodenum and jejunum of the mice were showed, the fluorescence intensity were analyzed by Fiji and showed in (P) and (Q), respectively. Scare bars, 100 μm, 20 μm for the enlarged images, \**p* < 0.05, \*\**p* < 0.01, \*\*\**p* < 0.001, \*\*\*\**p* < 0.0001.

### 2.5 Dynamin1 and Rab5 were required for polysaccharides uptake

Although the role of CLTC protein in polysaccharide absorption has been characterized in our above investigation, other proteins involved in polysaccharide uptake are largely unknown. Dynamin, best studied for its role in CME, is a prototypical member of a family of multi-domain GTPases involved in fission and remodeling of multiple organelles.[40] To examine whether dynamin plays a role in polysaccharide uptake, HIEC-6 cells were treated with dynasore (dynamin inhibitor) in presence of GFPBW1 and WGE. The results showed that the internalized GFPBW1 and WGE were substantially reduced after blockage of dynamin function (Figure 4E, F), suggesting the extracellular to intracellular transport of the polysaccharide is likely reliant on dynamin. Unexpectedly, uptake of GFPBW1 in IEC-6 cells was not affected by dynamin inhibition while the internalization of WGE and transferrin were sharply decreased after dynasore treatment (Figure 4G, H), indicating even uptake of the same polysaccharide is cell species-dependent. Since mammalian genomes contain three closely related tissue-specific dynamin genes, siRNAs against *DNM1* and *DNM2* were employed to determine which isoform was required for the polysaccharide uptake by the cells. Interestingly, uptake of GFPBW1 was impaired by si-*DNM1*, and the uptake of WGE and transferrin was almost completely abolished by *DNM1* depletion, while the knockdown efficiency of *DNM1* was confirmed by immunoblot (Figure 5D-F). Interestingly, although the expression of DNM2 was reduced sharply by si-*DNM2*, the internalization of GFPBW1 and WGE was not significantly disturbed by *DNM2* silencing (Figure S17, Supporting Information). Collectively, we speculated that DNM1 rather than DNM2 was responsible for polysaccharides uptake at least in the tested intestine cells. To further confirm this phenomenon *in vivo*, *DNM1* conventional knockout mice was generated firstly, unfortunately, we failed to obtain the homozygous *DNM1* KO mice due to the fact that homozygous *DNM1* offspring died after birth. Therefore, wild type (Ctrl) and heterozygous mice were employed to do the flowing exploration. Interestingly, both GFPBW1 and WGE were internalized in Ctrl mice intestine after polysaccharides orally administrated, while the intestinal internalized GFPBW1 and WGE were slightly or sharply decreased in *DNM1* KO heterozygous mice, respectively (Figure 5L, M). The above data suggested that dynamin 1 is required for the GFPBW1 and WGE internalized in the intestine cells.

Rab proteins are a large family of GTPases that regulate vesicle transport in cells. Rab5 is a marker of early endosomes and plays an important role in CME.[41] To understand the possible role of Rab5 in polysaccharide uptake mediated by clathrin, immunostaining with Rab5 was performed after both HIEC-6 and IEC-6 cells were incubated with GFPBW1 and WGE. Interestingly, Rab5 was partially or completely colocalized with the internalized GFPBW1 and WGE, suggesting that Rab5 probably involved in polysaccharides uptake (Figure 4H-K). Moreover, fluorescence imaging showed that the uptake of GFPBW1 and WGE was dramatically inhibited by *RAB5A* silencing, while Western blotting analyses confirmed the knockdown efficiency (Figure 5G-I), suggesting that Rab5 was also critical for polysaccharides uptake. To further evaluate the role of Rab5 in polysaccharide trafficking *in vivo*, *RAB5A* conventional knockout mice were generated. Then the fluorescein-labeled GFPBW1 or WGE were orally administrated to both wild type (Ctrl) and *RAB5A*^-/-^ (*RAB5A* KO) mice. Indeed, internalized polysaccharides in both duodenum and jejunum segments of the Ctrl mice were observed obviously, while trace amounts of internalized polysaccharides could be detected in *RAB5A* KO mice, indicating that littermates of the *RAB5A* KO mice displayed a marked impairment in their ability to internalize polysaccharides (Figure 5N-Q). Taken together, these data demonstrated that Rab5 was also required for polysaccharides uptake *in vitro* and *in vivo*.

### 2.6 Internalized polysaccharides were detectable in endosome, lysosome, Golgi and endoplasmic reticulum in intestinal cells

Further, we were wondering how the intracellular fates of the internalized polysaccharides are after the uptake? To address this question, HIEC-6 cells were incubated with fluorescein-labeled GFPBW1 and WGE for the indicated time and then immuno-stained with EEA1 (endosome marker), LAMP1 (lysosome marker), AIF (mitochondria marker), RCAS1 (Golgi marker) or PDI (endoplasmic reticulum marker), respectively. Both GFPBW1 and WGE were mainly colocalized with EEA1 as expected (Figure 6A, B). Furthermore, significant amount of colocalization between the internalized GFPBW1, WGE with LAMP1 was also observed (Figure 6C, D). However, it seems that both polysaccharides were not much in mitochondria, since there was little detectable association between polysaccharides and AIF (Figure 6E). Moreover, we found that the internalized polysaccharides were significantly colocalized with the marker of Golgi apparatus-RCAS1 (Figure 6F, G). Finally, the internalized polysaccharides were mainly colocalized with PDI, which is the marker of endoplasmic reticulum (ER) (Figure 6H, I). Thus, based on the above results, we speculated that polysaccharide might be dynamically transported among organelles. Upon internalization, polysaccharides could move from the early endosomes to lysosomes or/and further moved to other organelles such as Golgi, ER but not mitochondria.

**Figure 6.**
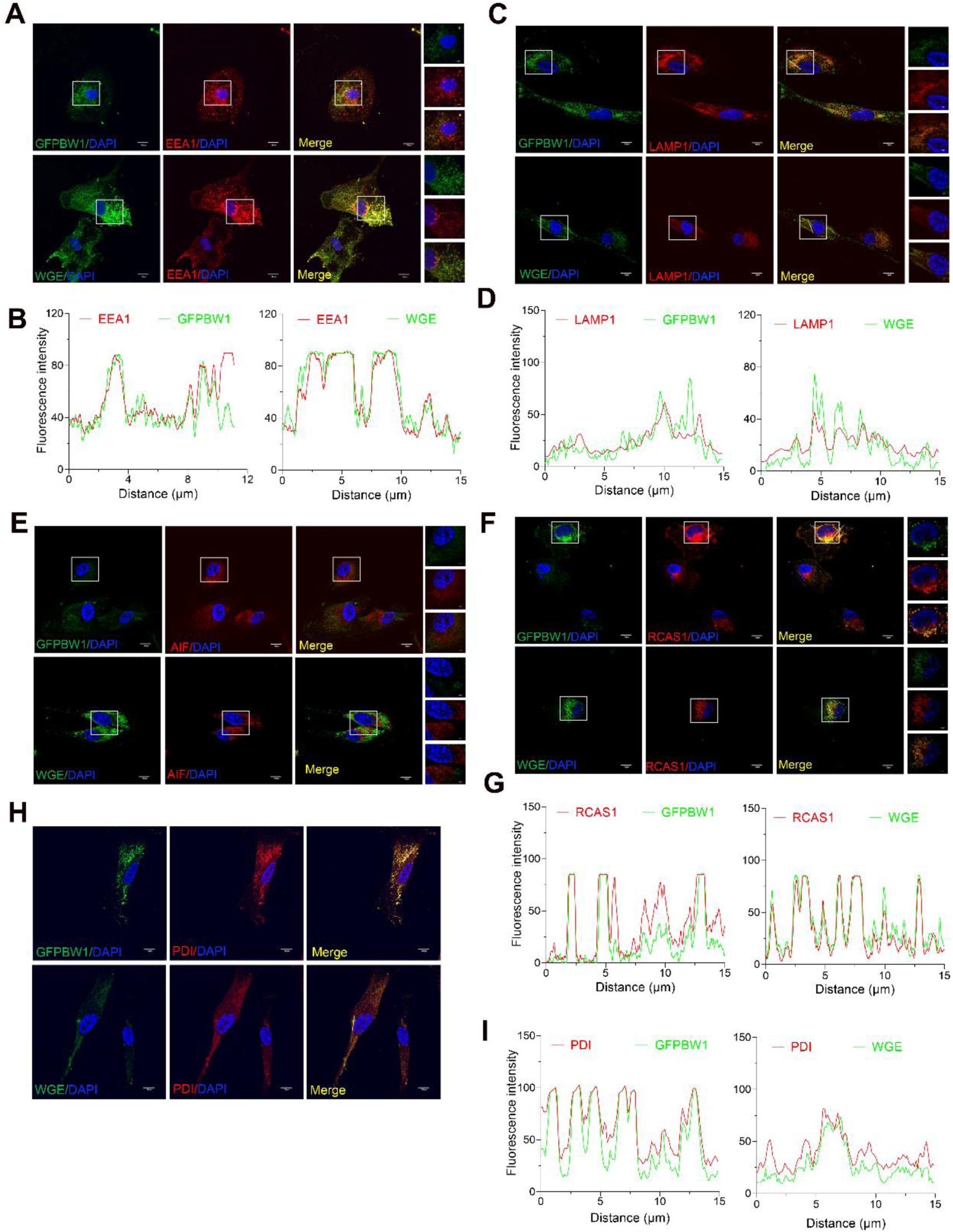
The internalized polysaccharides were located in endosome and lysosome, Golgi, endoplasmic reticulum (ER) but not in mitochondria in HIEC-6 cells. Cells were treated with labeled GFPBW1 and WGE for 1 h, then the cells were immune-stained with EEA1 (A), AIF (E), RCAS1 (F) and PDI (H), the colocalization analysis results were exhibited in (B), (G) and (I), respectively. Meanwhile, HIEC-6 cells were incubated with GPFPBW1 and WGE for 1.5 h and immune-stained with LAMP1 (C), the colocalization analysis are in (D). Scale bars, 10 μm, 2 μm for the enlarged images.

### 2.7 EGFR, Dectin-1 and BMPRIA were implicated in polysaccharides uptake

It is reported that transport of some polysaccharides was mediated by specific receptors.[42] Previously, we found the sulfation derivative of WGE could bind to BMPRIA while pectin RN1 (branched l,6-linked galactan with arabinose) could bind to EGFR,[43] indicating these receptors might mediate some specifical function of exopolysaccharides. Additionally, dectin-1 was the well-known receptor for β-1,3 glucan, this phenomenon is also demonstrated by our group evidenced by the binding between GFPBW1 and dectin-1 in our previous work.[32] To further understanding the role of cell membrane receptors in polysaccharides uptake, EGFR was firstly silenced in HIEC-6 cells in the presence of GFPBW1 and WGE. The results showed that the uptake of both polysaccharides was dramatically diminished, while the internalization of transferrin showed no difference between the control and EGFR knockdown cells, while the efficiency of siRNAs was confirmed by immuno-blotting (Figure 7A-C). Furthermore, we observed similar phenomenon in T24 cells either by Gefitinib (EGFR inhibitor) or by si-*EGFR* interference (Figure S18-S19, Supporting Information), further validated the important role of EGFR in polysaccharides uptake. Secondly, siRNAs were utilized to knockdown the expression of dectin-1 in HIEC-6 cells, the fluorescence images revealed that the internalization of GFPBW1 was significantly abolished, while the uptake of transferrin did not affect (Figure 7G-I). Meanwhile, we validated the importance of dectin-1 in GFPBW1 internalization using anti-dectin-1 treatment in T24 cells (Figure S20, Supporting Information). Thirdly, the internalization of WGE was indeed dramatically blocked by *BMPRIA* silence using siRNA in HIEC-6 cells, while transferrin showed no differences (Figure 7D-F). Moreover, the penetration of WGE was sharply blocked after BMPRIA depletion via anti-BMPRIA antibody treatment or siRNA interference in T24 cells (Figure S21-S22, Supporting Information). Hence, we speculated that the receptors EGFR, BMPRIA and dectin-1 was involved in the uptake for their corresponding polysaccharides.

**Figure 7.**
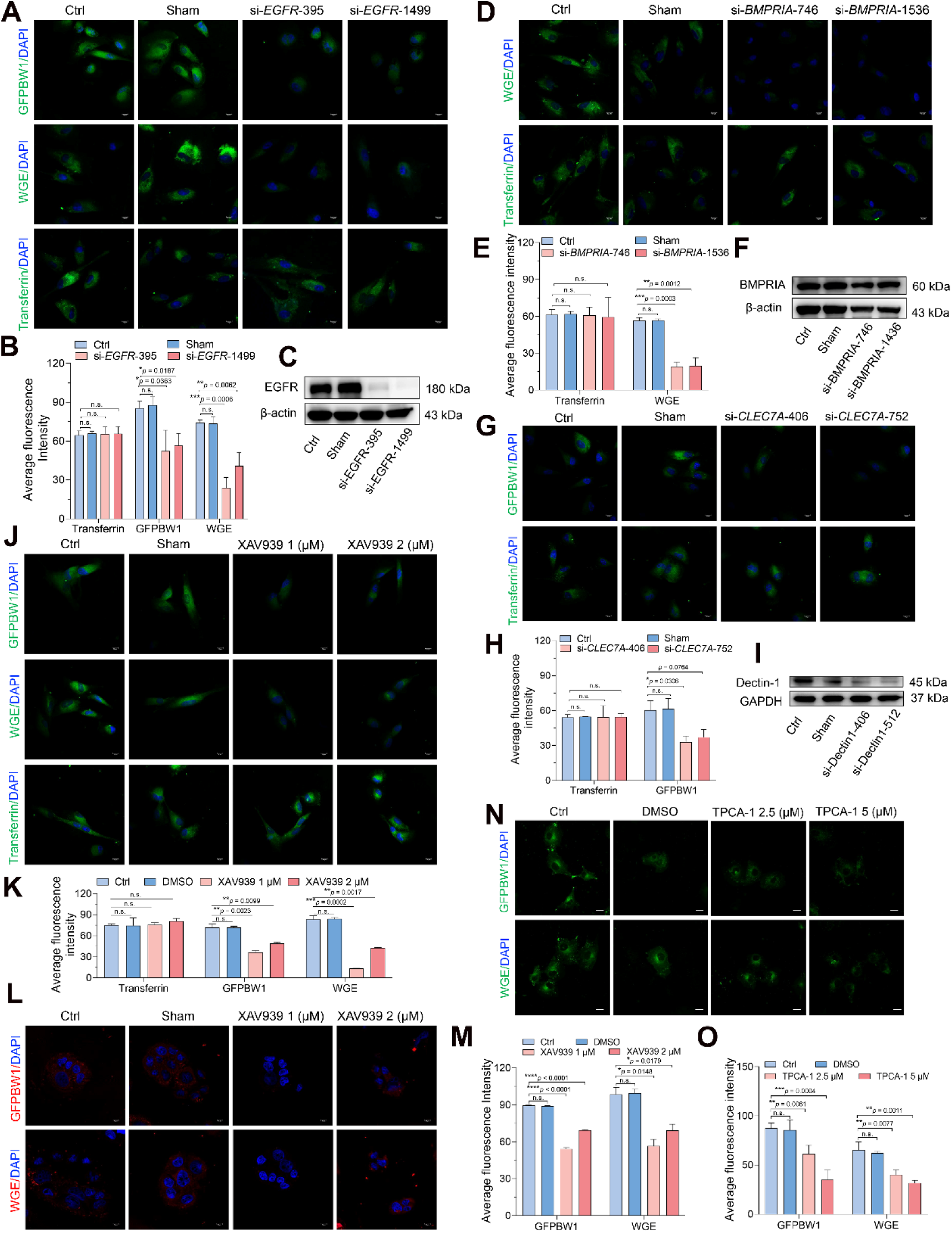
Blockage of EGFR, BMPRIA, dectin-1, wnt/β-catenin and NF-κB signaling pathway could suppress polysaccharide internalization in cells. The uptake results of GFPBW1 and WGE in HIEC-6 cells after transfecting with si-*EGFR* or si-*BMPRIA* or si-*CLEC7A* are displayed in (A), (D) and (G), the fluorescence intensity was quantified and are exhibited in (B), (E) and (H), respectively. Western blotting was used to detect the expression of EGFR (C), BMPRIA (F) and Dectin-1 (I) to confirm siRNAs efficiency. (J-M) HIEC-6 and Caco-2 cells were pretreated with XAV939 or DMSO, the images of the uptake experiments are showed in (J) and (L), the mean intensity was measured and exhibited in (K) and (M). (N) T24 cells were pretreated with TPCA-1, then the cells were incubated with GPFBW1, WGE and imaged, the intensity was quantified in (O). Scare bars, 10 μm. \**p* < 0.05, \*\**p* < 0.01, \*\*\**p* < 0.001, \*\*\*\**p* < 0.0001.

### 2.8 Wnt/β-catenin and NF-κB signaling pathways play a role in polysaccharides uptake

To further understand whether some signaling pathways are also implicated in the polysaccharide uptake, HIEC-6, Caco-2 or T24 cells were pretreated with XAV939 (Wnt/β-catenin signaling inhibitor) then incubated with fluorescein labeled GFPBW1 and WGE. The results showed that internalization of the two polysaccharides was significantly attenuated, indicating wnt/β-catenin signaling might be implicated in polysaccharides uptake (Figure 7J, M; Figure S23, Supporting Information). Interestingly, uptake of GFPBW1 and WGE was also sharply blocked by TPCA-1 (NF- κB inhibitor) (Figure 7N, O). Thus, these findings revealed that wnt/β-catenin and NF- κB signaling pathway might be critical in polysaccharide uptake.

### 2.9 Polysaccharide could be internalized in T24 cells via CME, dynamin1 and Rab5 was required during this process

To understand whether the phenomenon regarding the polysaccharide uptake is widely existed and to exclude individual cells variations and test the cell type specificity of polysaccharide uptake, a cancer cell line with epithelial morphology named as T24 was employed in this study. Firstly, the uptake dynamics of GFPBW1 and WGE were examined in T24 cells. The results demonstrated that uptake of GFPBW1 showed similar pattern in T24 cells with that in HIEC-6 cells among different time points, while WGE exhibited a different manner with that in intestine cells. Meanwhile, both GFPBW1 and WGE displayed a concentration-dependent manner as that in HIEC-6, IEC-6 and Caco-2 cells (Figure 8A-D).

**Figure 8.**
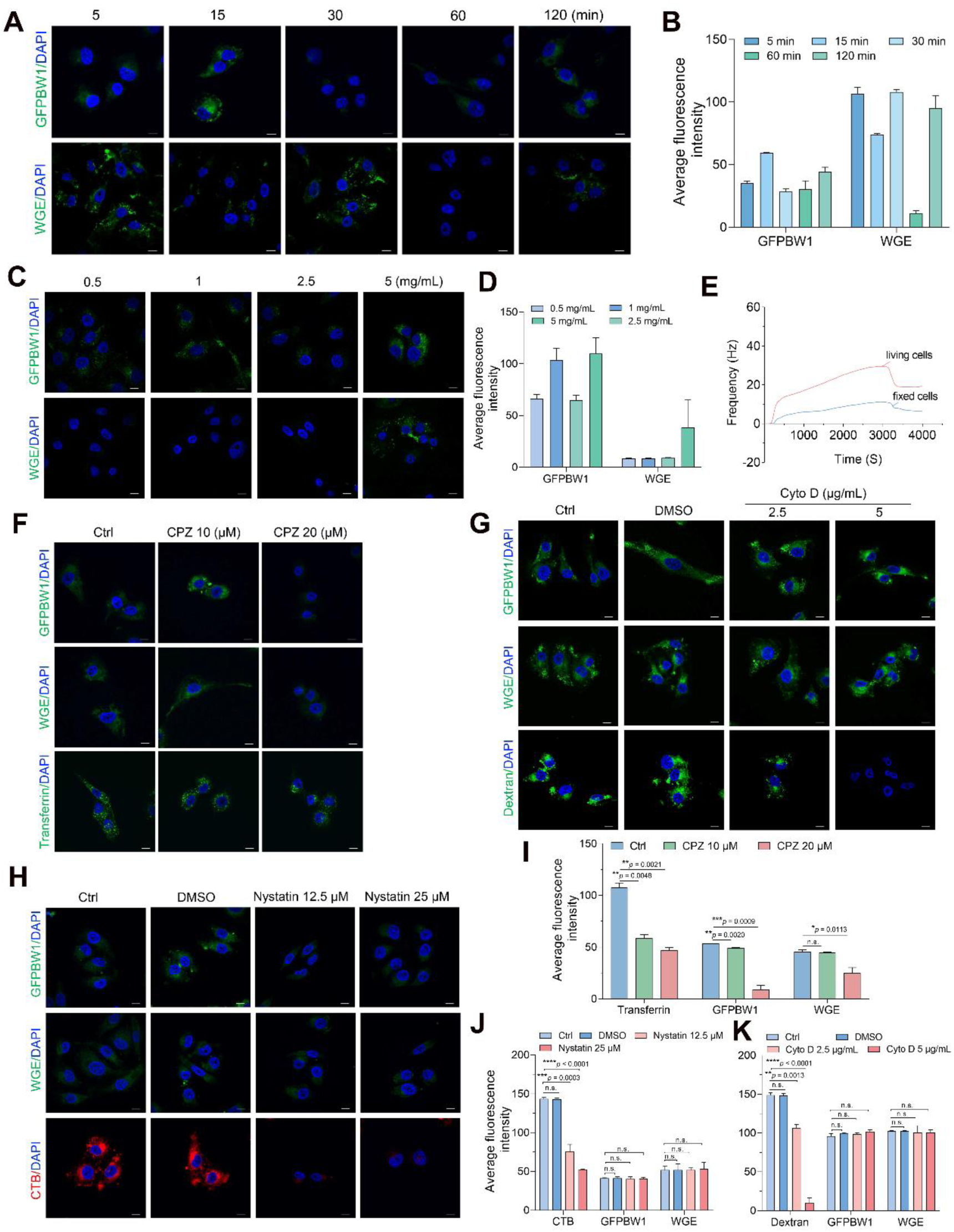
Polysaccharides could be internalized in T24 cells, and clathrin mediated endocytosis played an essential role during this process. (A-B) T24 cells were treated with fluorescein labeled GFPBW1 and WGE for the indicated time points, the images were detected by microscopy, the quantification results were shown in B. (C-D) The effects of polysaccharides concentration on their uptake in T24 cells were examined, and the images and quantification results were displayed in (C) and (D), respectively. (E) The binding curves for the binding of GFPBW1 to living and immobilized T24 cells detected by QCM are displayed. (F-H) T24 cells were pretreated with the inhibitors of various endocytosis routes for the indicated time intervals, then the uptake experiments of GFPBW1 and WGE were performed, the images are displayed in (F), (H) and (G), while the fluorescence intensity were quantified in (I), (J) and (K) separately, scale bars, 10 μm.

To gain insight to the dynamic uptake of polysaccharide, the interaction of GFPBW1 to living and immobilized T24 cells was detected by quartz crystal microbalance (QCM) biosensor which have proved to be a powerful tool to study macromolecular interactions.[44] Interestingly, the binding activity of GFPBW1 to living T24 cells was dramatically stronger than that of the immobilized cells, suggesting that GFPBW1 was dynamically internalized into living T24 cells (Figure 8E).

Next, similar with what we had done in intestine cells, inhibitors were employed to preliminarily determine the uptake route of polysaccharides in T24 cells. CPZ treatment caused a dramatic decrease in both uptake of GFPBW1 and WGE, while no obvious changes were observed in neither Nystatin nor Cyto D treated cell compared with Ctrl (Figure 8F-K), revealing that CME might be also critical in polysaccharides uptake in T24 cells.

To further confirm this result, the immunostaining of CLTC was conducted. Indeed, most of the internalized polysaccharides were colocalized with CLTC as expected (Figure S24, Supporting Information). Moreover, we found the binding of GFPBW1 to living T24 cells was blocked by anti-CLTC treatment, further confirmed the critical role of clathrin in GFPBW1 uptake (Figure S25A, Supporting Information). Subsequently, we found that knockdown *CLTC* with siRNA also cause internalization inhibition of both GFPBW1 and WGE, while the siRNA efficiency was checked by immunoblot in T24 cells (Figure S26, Supporting Information). Furthermore, we also utilized QCM to confirm siRNA efficiency. Interestingly, the binding curve of the Ctrl group was much higher than that of si-*CLTC* group after GFPBW1 treatment, indicting the inhibition of GFPBW1 internalization was caused by *CLTC* knockdown (Figure S25B, C, Supporting Information). To further confirm, we wondered whether overexpression of clathrin would promote polysaccharide internalization. To test, full-length *CLTC* cDNA plasmid was transfected in T24 cells, Western blotting analysis confirmed the over-expression of *CLTC*. However, the amount of internalized GFPBW1 or WGE was not significantly increased (Figure S27, Supporting Information). Interestingly, that polysaccharides could not colocalize with caveolin-1 in T24 cells, further confirmed that polysaccharides uptake did not require caveolin (Figure S28, Supporting Information).

Similar to our investigation in intestine cells, the immunostaining with Rab5 was performed and the results showed that Rab5 could colocalize with internalized GFPBW1 and WGE in T24 cells. Furthermore, the uptake of GFPBW1 and WGE was blocked after *RAB5A* depletion, while the knockdown efficiency was examined by immunoblot (Figures S29-S30, Supporting Information). Similarly, T24 cells were cultured with fluorescein labeled polysaccharides, followed by immuno-staining with anti-dynamin antibody. Indeed, the internalized GFPBW1 and WGE were colocalized with dynamin on the cell membrane or in cytoplasm, indicating that dynamin may also be involved in polysaccharides uptake. Subsequently, the expression of DNM1 was disturbed by siRNA, the microscopy analysis indicated that the uptake of GFPBW1 and WGE were sharply blocked in the *DNM1* silence T24 cells (Figure S31-S32, Supporting Information). Meanwhile, the internalization of the polysaccharides was dramatically attenuated by dynamin inhibition caused by dynasore. Similar to that of intestine cells, we also found that the internalization of polysaccharides did not affect by *DNM2* disturbing (Figure S33-S34, Supporting Information).

Eps15 (an acronym for EGFR pathway substrate 15) is an important protein involved in CME.[45–46] To determine the role of Eps15 in polysaccharides uptake, immunostaining assays were performed in IEC-6 and T24 cells after treatment with labeled polysaccharides. The results revealed that GFPBW1 and WGE were co- localized with Eps15 in most areas of the cytoplasm in both cells, suggesting that Eps15 might also involve in polysaccharide uptake (Figures S35-S36, Supporting Information).

### 2.9 The internalized polysaccharides might be examined in endosome, lysosome and Golgi in T24 cancer cells

The above results outlined extracellular to intracellular transport of polysaccharides in human intestinal epithelial cells, but how is their intracellular feature in cancer cells? Thus, similar strategy was employed as we used in HIEC-6 cells. The immuno-staining experiments revealed that both GFPBW1 and WGE showed strong colocalization with EEA1 and LAMP1 as expected (Figure S37-S38, Supporting Information). Moreover, we found the internalized polysaccharide were significantly colocalized with RCAS1, consistent with the results in HIEC-6 cells (Figure S39, Supporting Information). However, by using AIF and PDI as markers for mitochondria, ER, respectively, we found that there was no obvious association between polysaccharides and AIF or PDI (Figure S40-S41, Supporting Information). Thus, we speculated that polysaccharide could transport among organelles in cancer cells, they could move from the early endosomes to lysosomes and might further deliver to Golgi, but not mitochondria or ER in cancer cells.

This study demonstrated that some natural polysaccharides could be absorbed in intestine after oral administration via CME including dynamin1, Rab5 as the co-factor proteins based on the results of genetic, *in vivo*, *in vitro* and pharmacological experiments. The major findings of this study include the following: (i) Different linkage and various charged polysaccharide tested in this study were able to penetrate the Caco-2 monolayer with the *P*_app_ values of all the tested polysaccharides were greater than 1 × 10^-6^ cm/s. Therefore, theoretically, they could be absorbed *in vivo* according to the correlation between oral drug absorption and *P*_app_ values in Caco-2 model.[34] (ii) By using two different model polysaccharides: GFPBW1 (β-1,3 linked glucan, 300 kDa) and WGE (α-1,4 linked glucan, 700 kDa), we characterized *in vivo* absorption features in rats and mice using unlabeled, fluorescein-labeled, and radioactive (^99m^Tc, ^3^H) labeled strategy. Surprisingly, at least some branched β-1,3-andα-1,4 linked glucan might be detected in mice and rat plasma without significant degradation evidenced by little shift in their retention time on HPLC before and after the polysaccharide oral administration, suggesting that at least some polysaccharides might be absorbed as original macromolecular form with little degradation and could penetrate the intestinal lining and be transported to the blood circulation system. (iii) Although some researchers have tried to move the field forward by figuring out the pharmacokinetic characteristics of several natural polysaccharides,[18,19]the mechanism underlying polysaccharides internalization remained enigmatic. In the present study, we show that clathrin contributes to intracellular polysaccharide trafficking by endocytosis, a process also requiring dynamin 1 and Rab5 as well as Eps15 (Figures 4, 5 and Figures S35-S36, Supporting Information). Depletion of clathrin, Rab5 or dynamin 1 leads to sharp reduction in polysaccharide uptake in both *in vitro* and *in vivo* studies. To our knowledge, this is the first study to comprehensively explore polysaccharide uptake mechanism using pharmaceutic agents, siRNAs and knockout mouse models. (iv) Previous study regarding to natural polysaccharides uptake was mostly investigated in Caco-2 cells, which is actually colorectal cancer cell. In this study, we demonstrated the universality of polysaccharides internalization *in vitro* using normal human, rat intestinal cell line and cancer cell lines. We showed that polysaccharides could internalize into all the tested cell lines and the uptake dynamics is somehow cell line specifical. (v) We characterized the intracellular transport cellular destination of the internalized polysaccharides in intestinal cells and we also further showed that the critical roles of some specific receptors such as EGFR, BMPRIA and dectin-1 were somehow required for the tested polysaccharides uptake.

Most clinical biological products such as proteins and peptides are mainly delivered via parenteral route due to their negligible oral bioavailability attributing to the large molecular weight hampering the interaction with biological barriers, vulnerable to degradation by proteolytic enzymes, and their high hydrophilicity. Recently, extensive studies make efforts to develop alternative delivery strategies including permeation enhancers, enzyme inhibitors, and mucoadhesive polymers.[47] Lee group designed a hydrophobic ion-pairing (HIP) of insulin and found that the *P_app_* of insulin was about 6.36 folds enhanced in Caco-2 models.[48] Furthermore, the transport mechanism of the exogenous proteins was also attracted researchers’ attention. Wang et al. showed that inhibition of CME predominantly blocked the transportation of 55-100 kDa proteins from the hemolymph into the fat body verifying by RNAi.[49] Bagnat and his group demonstrated that lysosome rich enterocytes internalize dietary protein via receptor mediated and fluid phase endocytosis for intracellular digestion and trans-cellular transport in the immature gut of zebrafish and suckling mice.[50] Thus, the major molecular mechanism underlying protein transport was endocytosis which is similar as the result we demonstrated in some natural polysaccharides uptake in this study. However, some natural polysaccharides seem to be more stable in gastrointestinal tract than protein. Our data show that at least partial of the absorbed polysaccharide was exhibited as macromolecular form with little degradation in plasma (Figures 1, 2).

Furthermore, we were also able to rule out that some routine endocytosis pathways such as lipid raft mediated endocytosis, caveolin-mediated endocytosis or macropinocytosis did not involve in the internalization of polysaccharides in human intestinal cells (Figure 3). More interestingly, the intracellular transport of the internalized polysaccharides was traced using colocalization analysis. We uncovered that the internalized polysaccharide could transport to endosome, lysosome, Golgi as well as ER but not mitochondria in HIEC-6 cells (Figure 6). What surprising us is that the destination of intracellular transport of polysaccharides relay on cell type. The polysaccharides could be transported to endosome, lysosome and Golgi but not ER or mitochondria in T24 cells (Figures S37-S41, Supporting Information). Nevertheless, the detailed mechanism underlying this difference still need further exploration.

Except clathrin and its partner proteins, we showed that some particular cell membrane receptors also play a role in polysaccharides uptake. Moreover, we demonstrated that as receptors, BMPRIA and dectin-1 mediated the endocytosis of WGE and GFPBW1, respectively (Figure 7). Since polysaccharides with different structure feature may bind to their specific receptors on cell membrane, more binding test need to be explored to discover the exact receptors which mediate special polysaccharide endocytosis.

## 3. Conclusion

Collectively, this study clearly demonstrate that some natural polysaccharides can be absorbed by intestine and enter blood circulation system after oral administration via serials of *in vitro* and *in vivo* evidences. We also showed that polysaccharides internalization was via CME with the synergistic action of Rab5, dynamin1 and Eps15. Furthermore, the receptors BMPRIA, dectin-1 and EGFR, and wnt/β-catenin, NF-κB signaling were also critical for polysaccharide endocytosis. However, more efforts should be put on studying of precise mechanisms and the interaction with polysaccharides and the targeting proteins.

## 4. Experimental Section

A detail description of the methods is provided in the Supporting Information.

## Acknowledgements

We thank Professor Zhichao Pei, Yuxin Pei from Northwest A & F University, China, for their help in QCM analysis, Professor Yuan Wang, Fei Wu from Shanghai University of Traditional Chinese Medicine, China, for their help in fluorescein-labeled polysaccharides analysis. We express our sincere gratitude to INFINITUS Co., Ltd. (Guangzhou, China) for their support for our work. This work was supported by grants from National Natural Science Foundation of China (Grant No. 82374059, 32271332, 31870801, 82341097), Lingang Laboratory (Grant No. LG202101-01-02), National Key R&D Program of China (2022YFA1303802). This work was partially supported by High-level New R&D Institute (2019B090904008), and High-level Innovative Research Institute (2021B0909050003) from Department of Science and Technology of Guangdong Province. We also thank Zhongshan Municipal Bureau of Science and Technology for their funding support.

## Author Contributions

W.F.L., D.X.C., and Y.W. conceived the experiments. W.F.L. carried out most of the experiments. X.C., J. Y. and P. P. W. purified the polysaccharides. Y.D.Z., P.F.D., and Z.M.W. helped the radioactive polysaccharides analysis. H. C., X. X. D. and K.P.W. helped analyze the data. K.D. directed the overall project and supervised the whole project. All authors read and approved the manuscript.

## Data Availability Statement

The data that support the findings of this study are available from the corresponding author upon reasonable request.

---

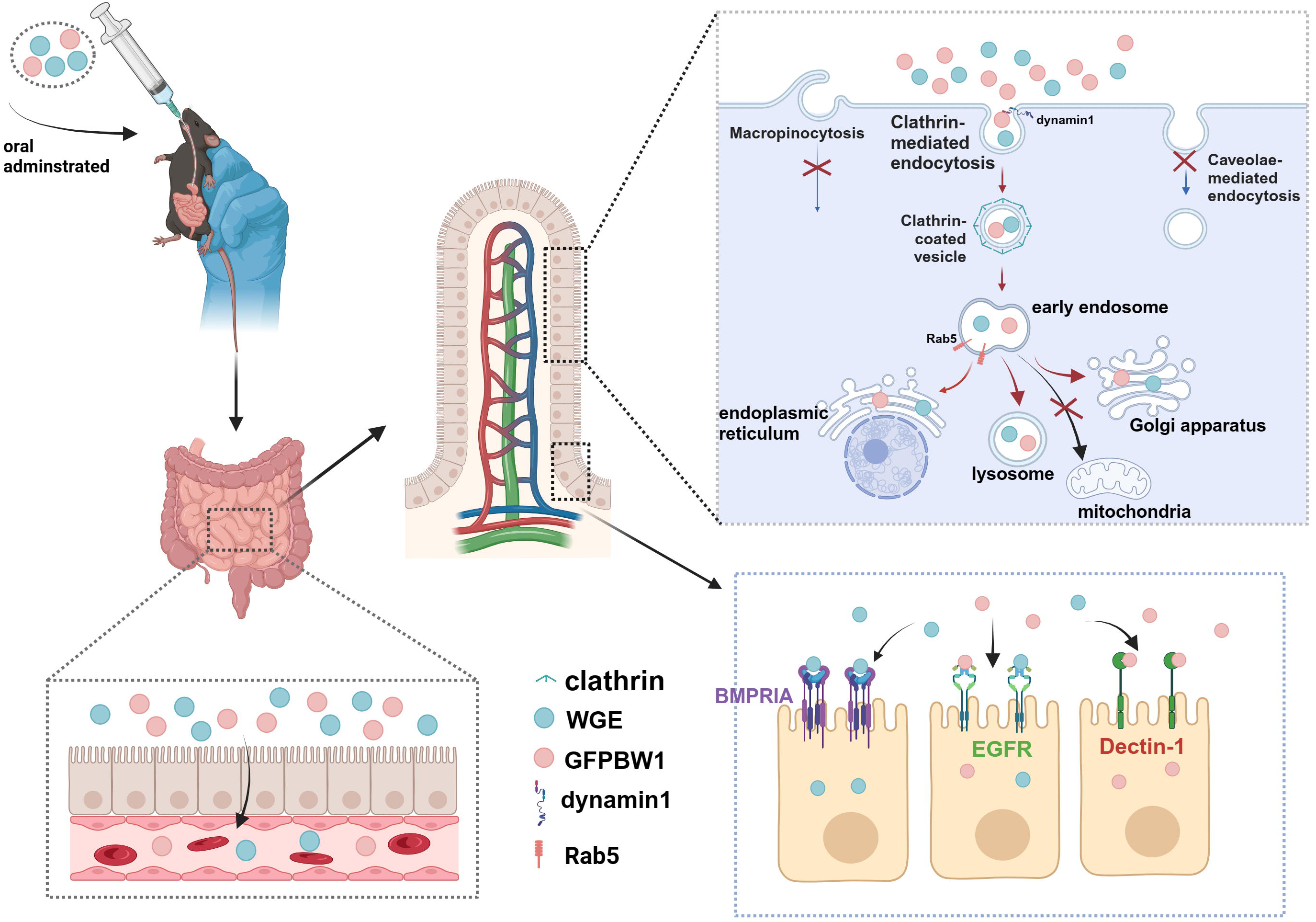

